# Two Regulators of G-protein signaling (RGS) proteins FlbA1 and FlbA2 differentially regulate fumonisin B1 biosynthesis in *Fusarium verticillioides*

**DOI:** 10.1101/2020.09.22.309260

**Authors:** Huijuan Yan, Zehua Zhou, Won Bo Shim

## Abstract

Fumonisins are a group of mycotoxins produced by maize pathogen *Fusarium verticillioides* that pose health concerns to humans and animals. Yet we still lack a clear understanding of the mechanism of fumonisins regulation during pathogenesis. The heterotrimeric G protein complex, which consists of Gα, Gβ, and Gγ subunits, plays an important role in transducing signals under environmental stress. Furthermore, regulators of G-protein signaling (RGS) proteins act as negative regulators in heterotrimeric G protein signaling. Earlier studies demonstrated that Gα and Gβ subunits are positive regulators of fumonisin B1 (FB1) biosynthesis and that two RGS genes, FvFlbA1 and FvFlbA2, were highly upregulated in Gβ deletion mutant ΔFvgbb1. *Saccharomyces cerevisiae* and *Aspergillus nidulans* contain a single copy of FlbA, but *F. verticillioides* has two putative FvFlbA paralogs, FvFlbA1 and FvFlbA2. Importantly, FvFlbA2 has a negative role in FB1 regulation. In this study, we further characterized functional roles of FvFlbA1 and FvFlbA2. While ΔFvflbA1 deletion mutant exhibited no significant defects, ΔFvflbA2 and ΔFvflbA2/A1 mutants showed thinner aerial hyphal growth while promoting FB1 production. FvFlbA2 is required for proper expression of key conidia regulation genes, including putative *FvBRLA, FvWETA*, and *FvABAA*, while suppressing *FUM21, FUM1*, and *FUM8* expression. Split luciferase assays suggest that FvFlbA paralogs interact with key heterotrimeric G protein components to impact *F. verticillioides* FB1 production and asexual development.

## 1. Introduction

*Fusarium verticillioides* (teleomorph: *Gibberella moniliformis* Wineland) is a fungal pathogen responsible for causing ear rot, stalk rot and seedling blight in maize worldwide. The fungus primarily utilizes conidia for dissemination, and the pathogen is capable of infecting and colonizing all developmental stages of maize plants (Blacutt, et al. 2018). Importantly, kernel infections by *F. verticillioides* lead to the production of fumonisins, a group of carcinogenic mycotoxins. Fumonisin B1 (FB1) is the most abundant and toxic form among fumonisin analogs, and long-term exposure to FB1 is linked to severe human and animal diseases including esophageal cancer and neural tube defects. Structurally, fumonisins contain a 19-20 carbon polyketide backbone, and usually multiple genes are involved in the biosynthesis of this complex group of secondary metabolites. A number of studies have demonstrated that genes involved in microbial secondary metabolite biosynthesis are organized as gene clusters (Alexander, et al. 2009, Kjaerbolling, et al. 2018, Ma, et al. 2010). The fumonisin biosynthesis gene cluster (referred to as the “*FUM* cluster”) was first discovered by Proctor et al (1999), which consists of 16 genes encoding biosynthetic enzymes and regulatory proteins (Proctor, et al. 2013). Inactivation of each of the key genes, e.g. *FUM1, FUM6, FUM8*, and *FUM21*, severely disturbed the production of fumonisins. Notably, previous studies demonstrated that these fumonisins-non-producing mutant strains did not significantly reduce maize ear rot in field tests (Desjardins, et al. 2002, Desjardins and Plattner 2000), which demonstrated that fumonisins are not essential for *Fusarium* ear rot pathogenicity. As we continue to seek strategies to minimize FB1 contamination in infested maize, we still lack a clear understanding on how *F. verticillioides* sense ambient environmental and host cues to regulate fumonisin biosynthesis.

G protein-coupled receptors (GPCRs) are the largest group of membrane receptors containing seven transmembranes (TMs) that transduce signals from the external environment to the cell, enabling the organism to adjust to its environment (Xue, et al. 2008). The canonical heterotrimeric G protein complex, which consists of α, β and γ subunits, plays important roles in transducing signals from GPCRs. When activated by specific ligands, GPCRs stimulate GDP to GTP exchange on the Gα subunit. Then, heterotrimeric G protein complex is dissociated into Gα subunit and Gβγ dimer, which triggers various downstream signaling pathways. GPCR signaling is known to be attenuated by G-protein-coupled receptor kinases (GRKs) and β-arrestin in animals, which are absent in filamentous fungi (Dohlman 2009). However, studies have shown regulators of G protein signaling (RGS) proteins in fungi act as GTPase-activating proteins, which promotes GTP hydrolysis of Gα subunit back to GDP-bound inactive form that terminates GPCR and G protein signaling pathways. RGS proteins typically contain 130-amino-acid RGS domains, which promotes the binding of RGS proteins to the Gα subunit. Notably, other than the RGS domain, RGS proteins are known to contain diverse non-RGS domains such as DEP (*Dishevelled, Egl-10* and *Pleckstrin*), PX, PXA, nexin C and TM, which are linked to various signaling pathways. For instance, the DEP domain in *Saccharomyces cerevisiae* ScSst2 was shown to interact with pheromone sensing Ste2 and mediated regulation of pheromone signaling responses (Ballon, et al. 2006).

*Aspergillus nidulans flbA* (for fluffy low brlA expression) was the first RGS protein identified in filamentous fungi, which is positively associated with conidiophore development and sterigmatocystin accumulation (Hicks, et al. 1997, Lee and Adams 1994). The *flbA* deletion mutant was not able to facilitate the transition from vegetative growth to conidiophore development. Conversely, overexpression of *flbA* led to premature *stcU* gene expressions and sterigmatocystin biosynthesis (Hicks, et al. 1997). In *Magnaporthe oryzae*, an AnFlbA1 ortholog MoRgs1 was shown to be involved in asexual development, pathogenicity, and thigmotropism (Liu, et al. 2007). Additionally, MoRgs1 showed physical interaction with a non-canonical GPCR MoPth11 and colocalized with Rab7, a late endosome marker (Ramanujam, et al. 2013). Another well studied RGS protein, MoRgs7 comprises of a N-terminal GPCR seven-transmembrane domain and a C-terminal RGS domain, which is critical for germ tube growth, cAMP signaling, and virulence in *M. oryzae* (Zhang, et al. 2011a).

Our earlier studies showed that functions of *F. verticillioides* Gβ and Gβ-like proteins are positively associated with FB1 production (Sagaram and Shim 2007, Yan and Shim 2020). Furthermore, transcription levels of four RGS genes *FLBA1, FLBA2, RGSB*, and *RGSC1* were significantly altered in Gβ deletion mutant ΔFvgbb1 when compared to the wild-type *F. verticillioides* strain (Mukherjee, et al. 2011). Unlike *A. nidulans, F. verticillioides* contains two putative FvFlbA paralogs, which were designated FvFlbA1 and FvFlbA2. Intriguingly, FvFlbA2 deletion mutation showed a drastic increase in FB1 production (Mukherjee, et al. 2011). Also, FlbA1 and FlbA2 were important for regulating host response during the fungal infection in surface-sterilized viable maize kernels. In this study, our aim was to further examine the regulatory mechanisms of the two FlbA genes in *F. verticillioides* and show how each gene plays unique roles and their relationship in FB1 biosynthesis.

## 2. Material and methods

### 2.1 Fungal strains and growth study

*F. verticillioides* M3125 was used as the wild-type strain in this study (Yan and Shim 2020). For growth and conidia production, all strains were grown on 0.2xPDA, myro, YEPD and V8 agar plates as described previously (Yan, et al. 2019). Spore germination assay was performed following our previously described methods (Yan and Shim 2020). For carbon utilization assay, Czapek-Dox agar was modified with various carbon sources such as sucrose (10g/L), dextrose (10g/L), fructose (10g/L) and xylose (10g/L) (Yan, et al. 2019). Fungal growth was determined by measuring colony diameter on agar plates after 8 days of incubation at room temperature. For mycelial weight assay, we inoculated 0.5 ml of WT and mutant conidia (10^6^/ml) into 100 ml YEPD broth, incubated in room temperature with constant shaking, and harvested at indicated time points. For stress assays, 4 μl spore suspension (10^6^/ml) was inoculated on 0.2xPDA agar plates and were grown with various stressors including 0.01% SDS, 2 mM H_2_O_2_, 0.6 M NaCl. All experiments were performed with at least three replicates. Inhibition rate was calculated as described previously (Yan and Shim 2020).

### 2.2 Gene deletion and complementation

Both ΔFvflbA1 and ΔFvflbA2 knockout mutants were generated in the wild-type strain via split-marker approach (Yan and Shim 2020). Briefly, partial hygromycin B phosphotransferase gene (*HPH*) designated as *PH* (929bp) and *HP* (765bp) were used to fuse with left and right flanking regions of the targeted gene with joint-PCR approach. All knockout constructs were amplified using Q5 High-Fidelity DNA Polymerase (New England Biolabs) except the second step of joint PCR using Taq enzyme (New England Biolabs) (Yu, et al. 2004). To further characterize the function of two FvFlbA paralogs in *F. verticillioides*, we generated the ΔFvflbA2/A1 double mutants in the ΔFvflbA2 background; partial geneticin resistance gene (*GEN*) designated as *GE* (1183 bp) and *EN* (1021 bp) were used to fuse with left and right flanking regions of *FvFLBA1* gene, and these constructs were transformed into with ΔFvflbA2 protoplast. Complementation fragments were amplified by Phusion Flash High-Fidelity PCR Master Mix (Thermo Scientific) and transformed into designated mutant protoplasts. All transformants were screened by PCR using Phire Plant Direct PCR Kit (Thermo Scientific) and verified by qPCR (Thermo Scientific). All primers are listed in Table 1.

### 2.3 Fumonisin B1, virulence, and gene expression assays

To study the FB1 production, cracked autoclaved kernels (2 g) and four surface sterilized kernels were used as described previously (Christensen, et al. 2012, Yan and Shim 2020). Briefly, once kernels were prepared in scintillation vials, fungal spore solutions (200 μL, 10^6^/mL) were inoculated in each vial and cultivated at room temperature for eight days. FB1 and ergosterol extraction and HPLC analyses were performed as described previously (Christensen, et al. 2012, Shim and Woloshuk 1999). FB1 levels were normalized to ergosterol contents. These experiments were performed with three biological replicates. Seedling rot was assayed by inoculating fungal spore suspension (5 μL, 10^6^/mL) and imaged after one-week growth in the dark room as described previously (Yan, et al. 2019). For qPCR analysis of conidiation-related genes and key *FUM* genes, mycelia were harvested from 7-day-old culture grown in myro liquid medium at 150 rpm. Primers for three conidia related genes, as well as *FUM1, FUM8* and *FUM21*, were described in previous studies (Yan, et al. 2019, Zhang, et al. 2011b). Relative expression levels of each gene were calculated using a 2^-ΔΔCT^ method and normalized with *F. verticillioides* β-tubulin gene (FVEG_04081). All qPCR assays were performed with three replicates.

### 2.4 Split luciferase complementation activity analysis

The coding sequences of PCR products were amplified by Q5 High-Fidelity DNA Polymerase and introduced to pFNLucG or pFCLucH by Gibson Assembly (New England Biolabs) (Kim, et al. 2012). Specifically, cDNAs of *FvFLBA1* and *FvFLBA2* were cloned into pFNLucH vector. cDNAs of *FvRAB5* (FVEG_00504), *FvRAB7* (FVEG_04809), *FvRAB11* (FVEG_11336), *FvVPS36* (FVEG_06233), *FvTLG2* (FVEG_07363), *FvPEP12* (FVEG_11540) were cloned into pFCLucH vector. *CLuc-FvGPA1, CLuc-FvGPA2, CLuc-FvGPA3, CLuc-FvGPB1*, and *CLuc-FvGBB1* were from our previous study (Yan and Shim 2020). Transformation, selection and luciferase activity determination were performed as described previously (Zhang, et al. 2018).

## 3. Results

### 3.1 Sequence analyses of FlbA1 and FlbA2 in *F. verticillioides*

Our previous study identified two FvFlbA paralogs in *F. verticillioides* including FvFlbA1 (FVEG_08855) and FvFlbA2 (FVEG_06192) (Mukherjee, et al. 2011). The new annotation in FungiDB (www.fungiDB.org) predicted that FvFlbA1 and FvFlbA2 encode a 499-amino-acid and 517-amino-acid protein, respectively. FvFlbA1 and FvFlbA2 share 52% identity at the amino-acid level. To identify FlbA orthologs in other fungi, *F. verticillioides* FvFlbA2 protein sequence was used in a BlastP query. Among fungal species we searched, only *Fusarium* species showed multiple copies of FlbA homologs. Both *F. verticillioides* and *F. graminearum* have two FlbA genes, and surprisingly, *F. oxysporum* has five putative FlbA paralogs. All FlbA orthologs are predicted to contain both DEP and RGS domains except that FvFlbA2 does not contain the RGS domain in the new annotation. Notably, FvFlbA1 and FvFlbA2 belong to different branches in our phylogenetic tree analysis (Fig. 1), which suggested that they may have diverged early and evolved to perform distinct functions in *F. verticillioides*. Phylogenetic tree and domain analysis followed a previous description (Yan and Shim 2020).

**Fig. 1.**
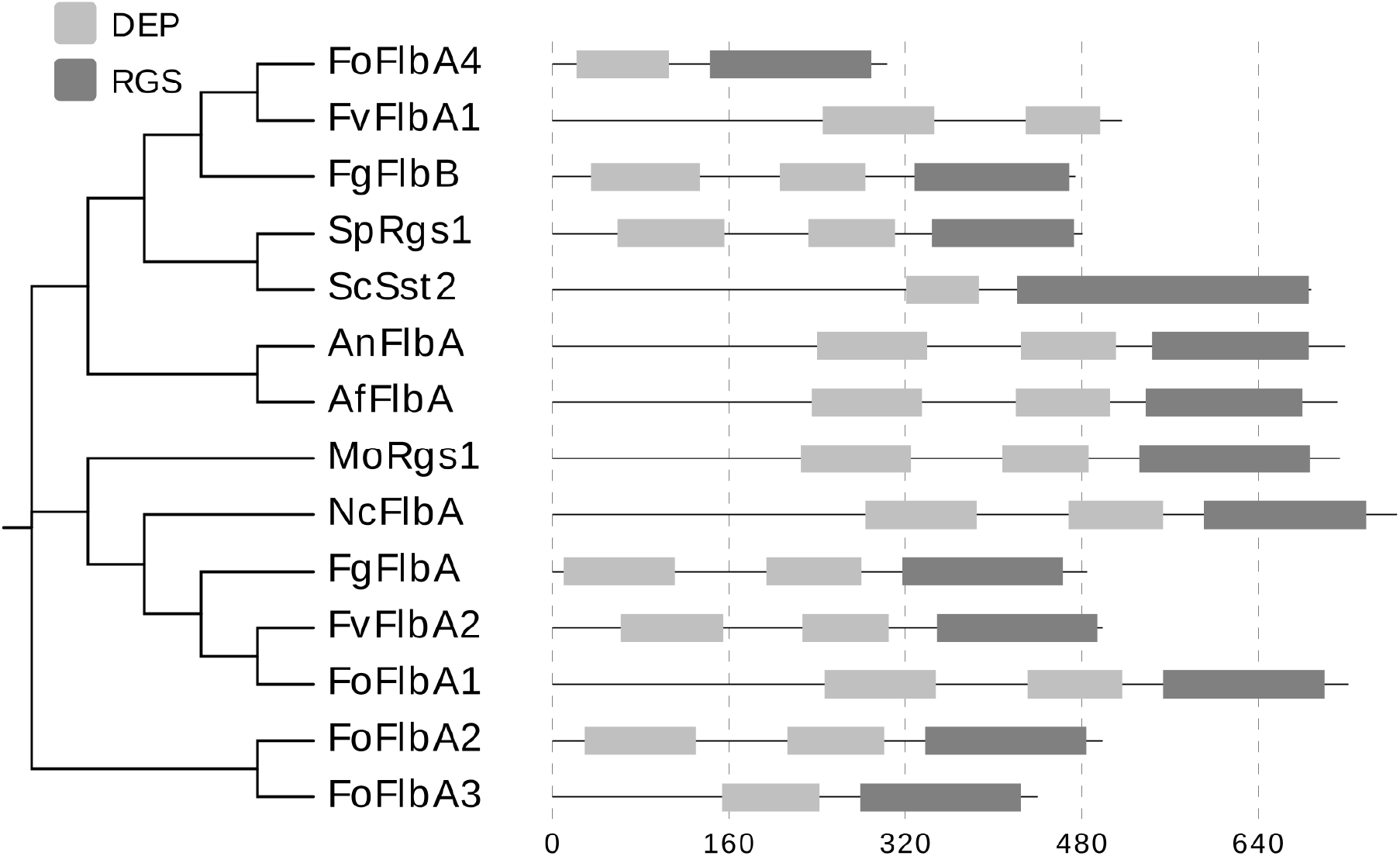
Phylogenetic and domain analysis of FlbA proteins in select fungal species. Organism names and NCBI locus tag: FvFlbA2 (FVEG_06192, 100% identify), FvFlbA1 (FVEG_08855, 52% identify), FoFlbA1 (FOXG_08482, 98% identity), FoFlbA2 (FOXG_06495, 74% identity), FoFlbA3 (FOXG_17640, 64% identity), FoFlbA4 (FOXG_09613, 53% identity), FoFlbA5 (FOXG_07099, 53% identity) in *F. oxysporum f. sp. lycopersici 4287*, FgFlbA (FGSG_06228, 98% identity), FgFlbB (FGSG_03597, 50% identity) in *F. graminearum*, MoRgs1 in *Magnaporthe oryzae* (MGG_14517, 63% identity), NcFlbA in *Neurospora crassa* OR74A (NCU08319, 74% identity), AnFlbA in *Aspergillus nidulans* FGSC A4 (An5893, 66% identity), AfFlbA in *A. fumigatus* Af293 (AFUA_2G11180,77%), SpRgs1 in *Schizosaccharomyces pombe* (SPAC22F3.12c, 31% identity), ScSst2 in *Saccharomyces cerevisiae S288C* (YLR452C, 36% identity). FoFlbA5 was not included in this phylogenetic analysis.

### 3.2 ΔFvflbA1 and ΔFvflbA2 mutants exhibit limited defects in vegetative growth

The two FvFlbA gene knockout mutants were generated by the split-marker approach (Fig. S2A). To confirm null mutations, *FvFLBA1* and *FvFLBA2* transcription levels in the mutants were tested by qPCR where expression of these target genes was not detectable when compared to the wild-type progenitor (Fig. S2D). The mutant ΔFvflbA1 showed no observable defect in terms of mycelial growth and conidiation on three different agar media. However, ΔFvflbA2 mutant showed slower growth on these media and also produced less conidia when compared to the wild type (Fig. 2). To further understand the relationship of two FvFlbA paralogs in *F. verticillioides*, we generated a ΔFvflbA2/A1 double mutant. ΔFvflbA2/A1 exhibited more severe defects in aerial hyphae (Fig. 2). In particular, the double mutant showed almost non-detectable level of conidia production when compared to the wild type and single gene mutants (Fig. 3A). Surprisingly, we discovered that ΔFvflbA2/A1 conidia, albeit the lower number of production, showed precocious conidial germination which was not observed in wild type or other mutants (Fig. 3B). Gene complementation strain ΔFvflbA2-Com demonstrated full recovery of growth defects observed in ΔFvflbA2 strain.

**Fig. 2.**
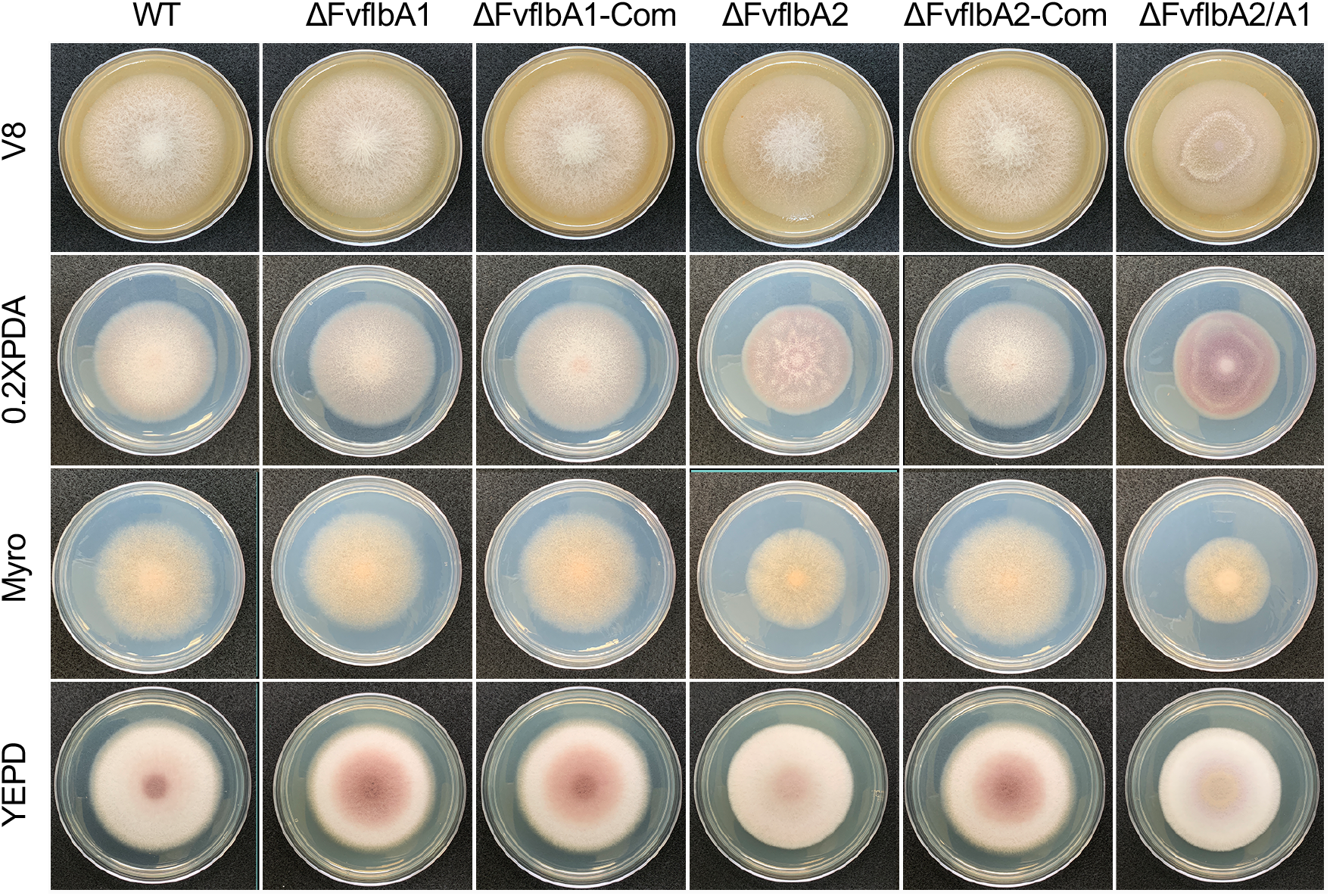
Hyphal growth of ΔFvflbA1, ΔFvflbA1-Com, ΔFvflbA2, ΔFvflbA2-Com, ΔFvflbA2/A1 strains. Colonies of the wild-type (WT), ΔFvflbA1, ΔFvflbA1-Com, ΔFvflbA2, ΔFvflbA2-Com, ΔFvflbA2/A1 strains were incubated on V8, 0.2xPDA, myro and YEPD agar at room temperature for 8 days.

**Fig. 3.**
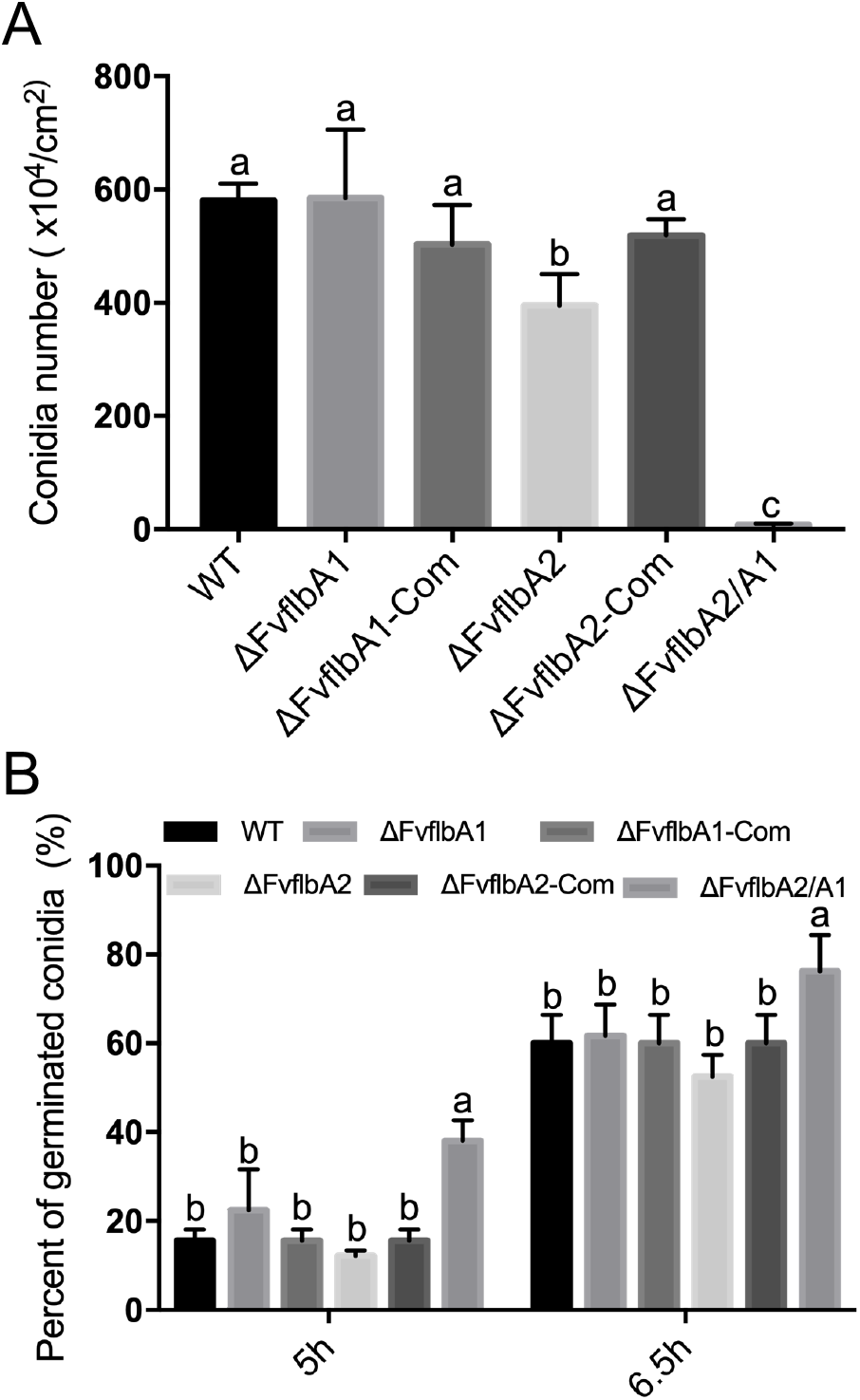
FvFlbA2 impacts on conidiation and germination. (A) Conidia were harvested from V8 agar plates after 8-day-incubation at room temperature. (B) WT, ΔFvflbA1, ΔFvflbA1-Com, ΔFvflbA2, ΔFvflbA2-Com, ΔFvflbA2/A1 strains were cultured in 0.2xPDB liquid medium with gentle shaking. Conidial germination rate was counted under a microscope. All experiments were performed with at least three biological replicates. The letters suggest statistically significant differences analyzed by Ordinary One-way ANOVA Fisher’s LSD test (p < 0.05).

### 3.3 FvFlbA2 is negatively associated with FB1 production

Our previous studies showed that RGS protein FvFlbA2 is negatively associated with FB1 production, while Gβ and one of the Gα proteins positively regulate FB1 biosynthesis (Mukherjee, et al. 2011, Yan and Shim 2020). To further understand the impact and relationship of the two FvFlbA genes in FB1 production, we analyzed FB1 levels by inoculating mutant strains on autoclaved cracked kernels and surface-sterilized living kernels (Fig. 4A and B). The results showed that ΔFvflbA2 and ΔFvflbA2/A1 produced significantly higher levels of FB1 compared with wild-type and ΔFvflbA1 strains (Fig. 4C and D). Surprisingly, ΔFvflbA2/A1 produced a drastically higher FB1 level (>300%) in contrast to ΔFvflbA2 strain when FB1 assay was performed using surface-sterilized maize kernels (Fig. 4D). Meanwhile, ΔFvflbA1 produced similar levels of FB1 as the wild-type and complemented strains. This result suggested that FvFlbA2 serves as a negative regulator of FB1 production, but the impact was more dramatic with double deletion of two FvFlbA paralogs and when viable host factors are associated with triggering FB1 biosynthesis.

**Fig. 4.**
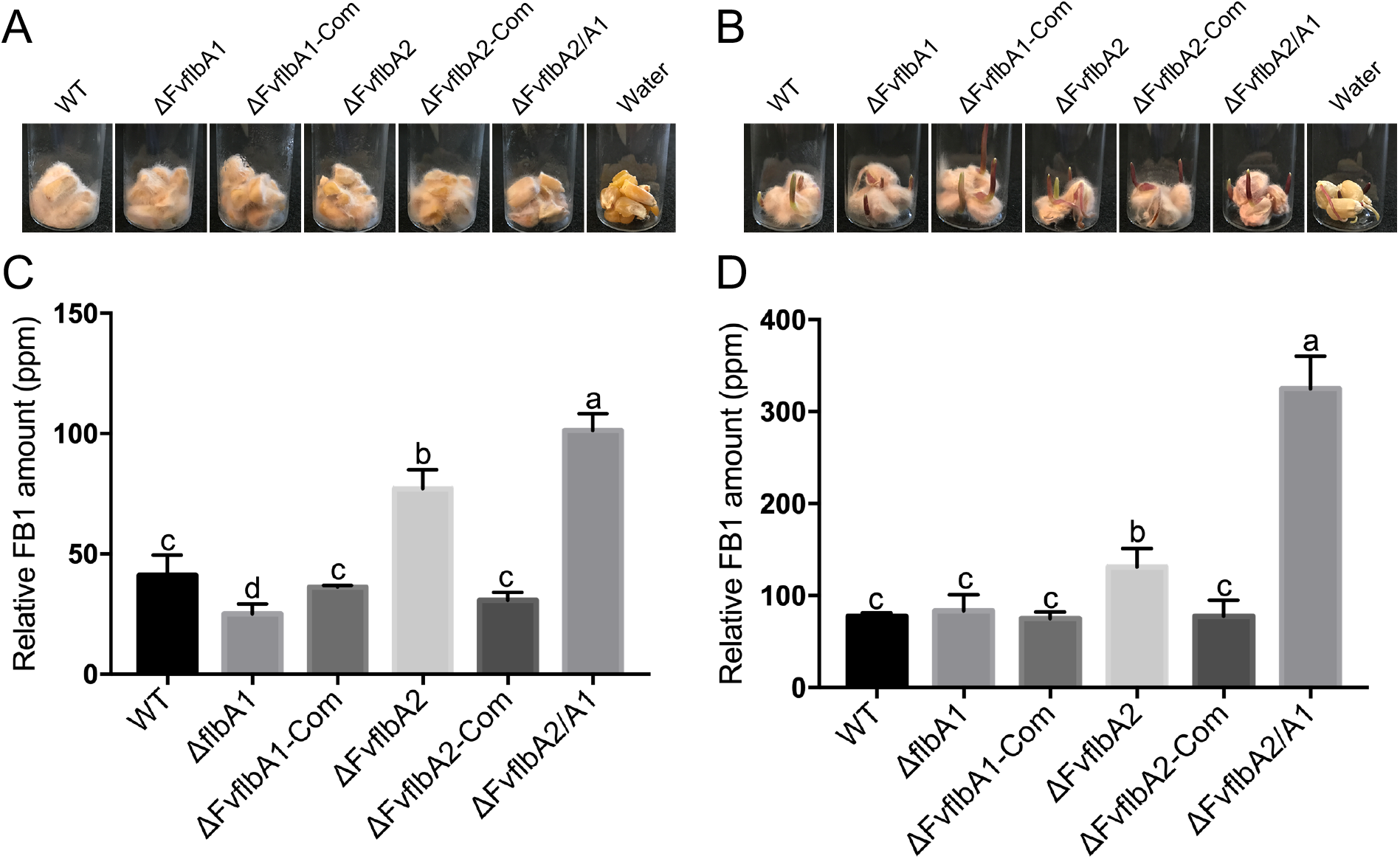
Colonization of two FvFlbA deletion mutants in kernels and FB1 assay. *F. verticillioides* wild-type (WT), ΔFvflbA1, ΔFvflbA1, ΔFvflbA2, ΔFvflbA2/A1 and complemented strains were cultivated in (A) 2-g cracked autoclaved kernels and (B) four surface-sterilized kernels for seven days at room temperature. (C) FB1 production in WT, mutants and complemented strains cultured on nonviable kernels and (D) viable kernels after seven days incubation at room temperature. The relative FB1 production levels were normalized to fungal ergosterol. All experiments were performed with at least three biological replicates. The letters suggest statistically significant differences analyzed by Ordinary One-way ANOVA Fisher’s LSD test (p < 0.05).

### 3.4 Expression levels of genes associated with conidiation and FB1 biosynthesis

The expression of *brlA* mRNA was not detectable in *A. nidulans flbA* mutant, which indicated that *flbA* is indispensable for *brlA* gene activation (Lee and Adams 1994). In our study, ΔFvflbA2 produced a lower count of conidia and germinated atypically in water in comparison to the wild type. To understand the impact of FvFlbA2 on asexual development at the molecular level, we analyzed the transcriptional expression of putative *BRLA*, *ABAA* and *WETA* genes in *F. verticillioides* wild-type and mutant strains. Our results showed that these three conidia-related genes were highly down-regulated in ΔFvflbA2 and ΔFvflbA2/A1 but not in ΔFvflbA1 (Fig. 5A). This result partially explains why FvFlbA2 function did not completely correlate with typical conidia production levels seen in *F. verticillioides*. To verify the role of the two paralogs in FB1 biosynthesis, we carried out qPCR analysis of three key *FUM* genes *FUM1, FUM8* and *FUM21*. Consistent with our FB1 results in myro liquid cultures (Fig. S3), key *FUM* genes were significantly up-regulated in ΔFvflbA2 and ΔFvflbA2/A1 (Fig. 5B). Importantly, our results suggested that FvFlbA1 and FvFlbA2 regulate distinct signaling pathways, but with FlbA2 playing a critical role in FB1 biosynthesis regulation.

**Fig. 5.**
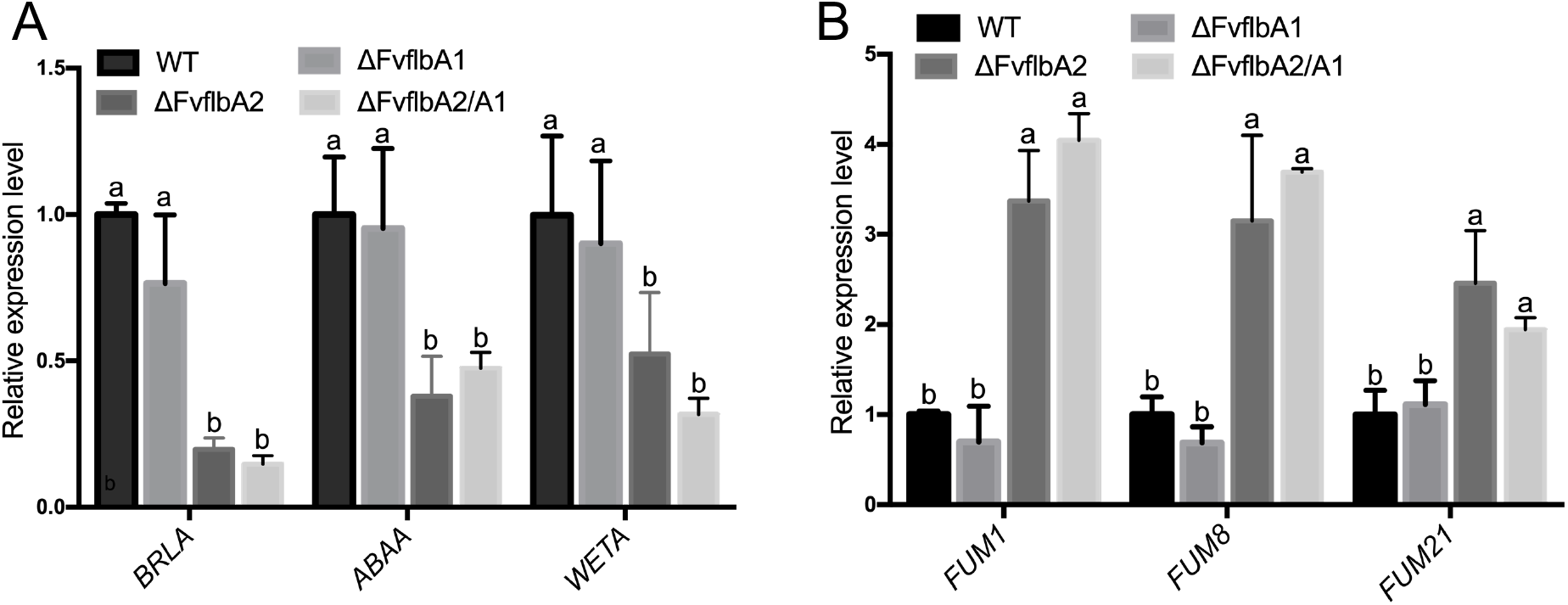
Effects of FvFlbA1 and FvFlbA2 on conidia-related genes and key *FUM* genes transcription. (A) Relative expression levels of putative *FvBRLA* (FVEG_09661), *FvABAA* (FVEG_00646) and *FvWETA* (FVEG_02891) in wild-type, ΔFvflbA1, ΔFvflbA2, ΔFvflbA2/A1 strains were normalized to *F. verticillioides* β-tubulin gene (FVEG_04081). (B) Relative mRNA expression level of three key *FUM* genes in deletion mutant strains in contrast to wild type. All experiments were performed with at least three biological replicates. The letters suggest statistically significant differences analyzed by Ordinary One-way ANOVA Fisher’s LSD test (p < 0.05).

### 3.5 *F. verticillioides* two FlbA paralogs are not required for stress responses, carbon utilization and virulence

To elucidate the roles of two FvFlbA proteins in response to environmental cues, we used various carbon sources including sucrose, dextrose, fructose and xylose to test if there are deficiencies in carbon sensing and utilization. Only minor vegetative growth defects were observed in ΔFvflbA2 and ΔFvflvA2/A1 mutants, suggesting that two FvFlbA paralogs are not critical for utilization of different carbon nutrients (Fig. 6A). Additionally, to test whether FvFlbA1 and FvFlbA2 are involved in cell wall integrity and various stress response signaling, we investigated the vegetative growth of mutants in the presence of SDS, H_2_O_2_, and osmotic (NaCl) stress agents in Czapek-Dox agar. We found that ΔFvflbA2 and ΔFvflvA2/A1 mutants exhibited trivial defects in growth in all media tested compared to the wild-type progenitor (Fig. 6B). However, ΔFvflbA2 and ΔFvflvA2/A1 mutants showed no significant differences in terms of inhibition rate compared to regular Czapek-Dox medium growth. Lastly, we found no difference in wild type and all mutant strains when tested for capacities to cause seedling rot (Fig. 7A and B). These results suggest that both FvFlbA1 and FvFlbA2 are dispensable for stress response, carbon utilization and virulence in *F. verticillioides*.

**Fig. 6.**
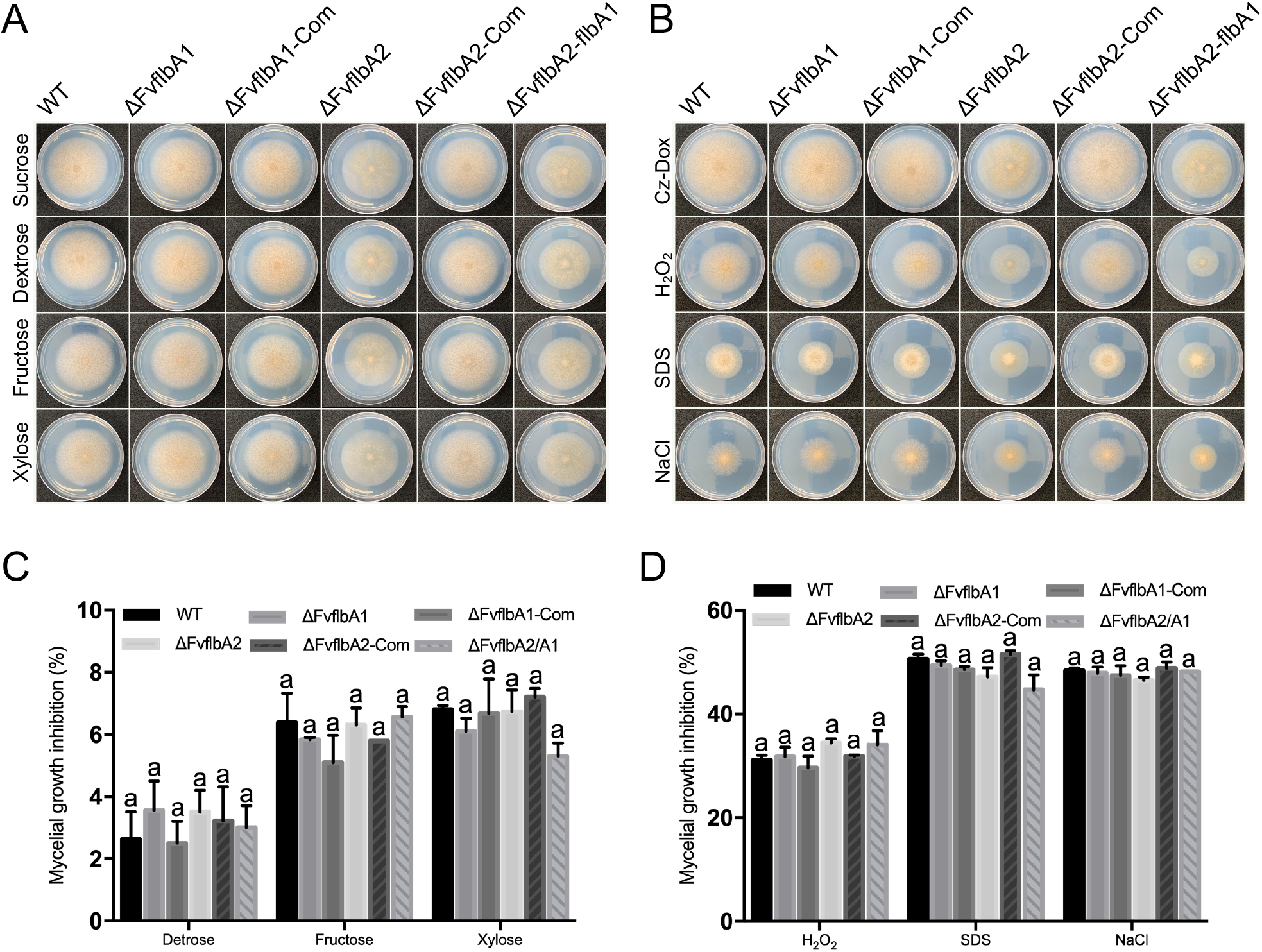
The influence of FvFlbA1 and FvFlbA2 on carbon utilization and stress response. (A) The colony of wild-type, ΔFvflbA1, ΔFvflbA2, ΔFvflbA2/A1 and complemented strains grown on modified Czapek-Dox agar plates with different carbon sources for 8 days. (B) Strains were cultured on Czapek-Dox agar plates with various stress agents for 8 days. The letters suggest statistically significant differences analyzed by Ordinary One-way ANOVA Fisher’s LSD test (p < 0.05).

**Fig. 7.**
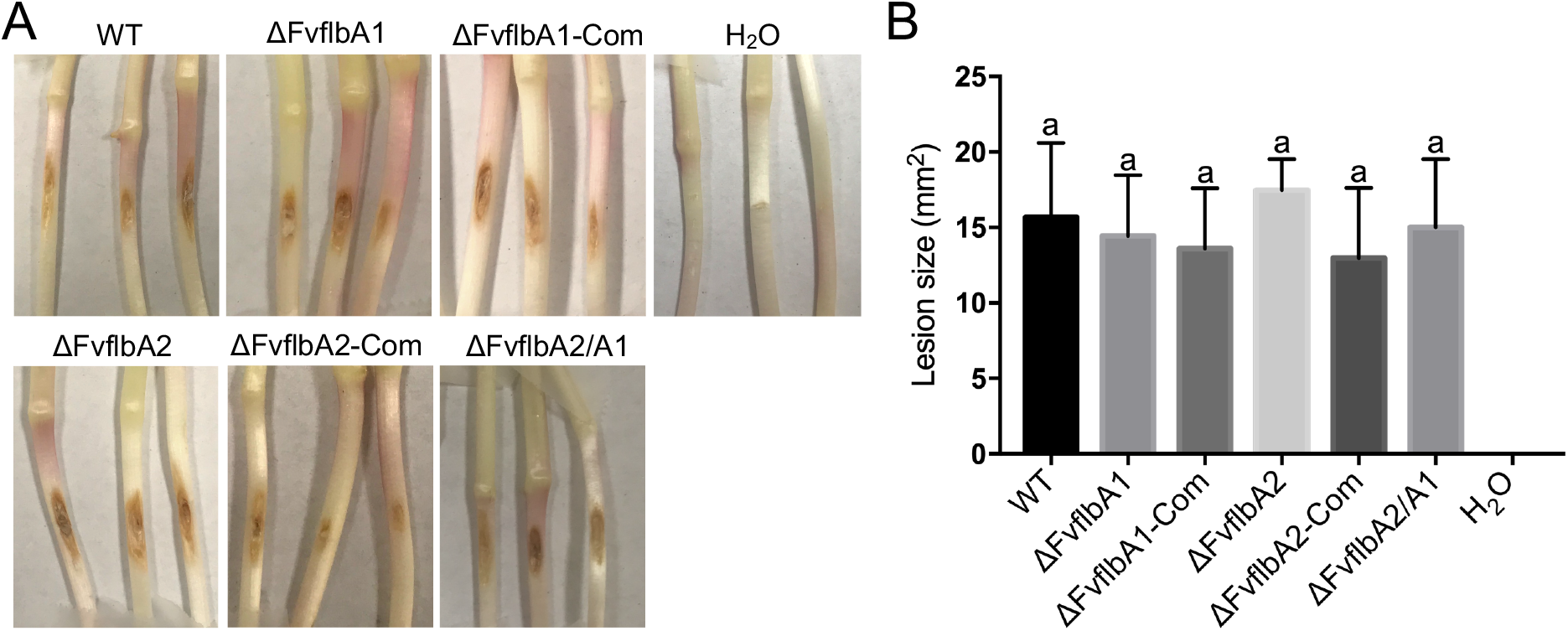
The pathogenicity of FvFlbA mutant strains in corn seedling rot assay. (A) A syringe needle was used to created wounds on one-week old silver queen seedlings. Spore suspensions (5 μl, 10^6^/ml) were inoculated on the wound sites. (B) The lesion size was quantified by Image J software after one-week incubation. All experiments were performed with at least three biological replicates.

### 3.6 FvFlbA1 physically interacts with FvFlbA2 and heterotrimeric G proteins components

The defects in growth and higher accumulation of FB1 production observed in ΔFvflvA2/A1 mutant raised a question whether FvFlbA1 and FvFlbA2 regulate other G protein signaling components in *F. verticillioides* through direct interaction. To test this, we performed split luciferase complementation assay in *F. verticillioides in vivo* (Kim, et al. 2012). We first tested FvFlbA1-NLuc interaction with FvFlbA2, canonical G protein components, as well as other proteins that are known to interact with RGS proteins. For instance, *S. cerevisiae* RGS protein Sst1 N-terminus showed interaction with multiple proteins involved in stress response signaling (ScVps36, ScTlg2, and ScPep12) and *M. oryzae* RGS protein MoRgs1 showing colocalization with a late endosome marker MoRab7 (Burchett, et al. 2002, Ramanujam, et al. 2013). Figure 8 shows FvFlbA1 interacting with all selected proteins in *F. verticillioides*, with particularly strong associations with FvGpa1, FvGbb1 and FvRab11. Luciferase activity was detected with FvVps36, FvTlg2 and FvPep12, but these were not strong. CLuc-FvFlbA2 showed interaction with FvFlbA1-NLuc. When we made FvFlbA2-NLuc to test interaction with these set of proteins, we did not observe luciferase activity. However, we acknowledge that this outcome must be further verified by other methods before concluding that FvFlbA2 does not directly associate with G protein signaling components in *F. verticillioides*.

**Fig. 8.**
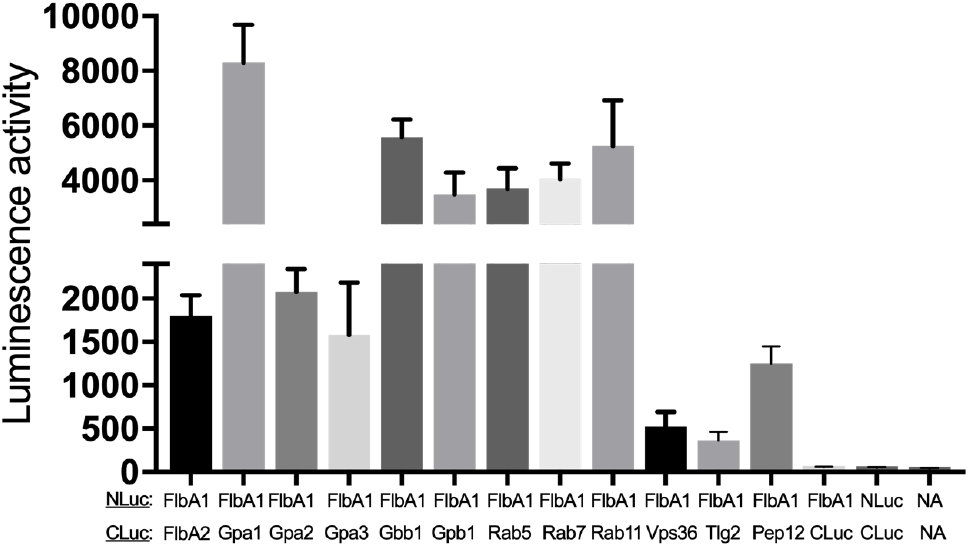
Interaction study between FvFlbA1 and FvFlbA1 in *F. verticillioides*. We used a split luciferase complementation assay to study the luminescence activity between FvFlbA1 and FvFlb2. Additionally, FvFlbA1 also exhibited significantly luminescence activity with FvVps36, FvTlg2 and FvPep12, FvRab5, FvRab7, and FvRab11 compared to negative controls. Negative controls included FvFlbA1 + CLuc, NLuc + CLuc, and no vector (NA) + (NA). Luminescence activity was gained from three replicates.

## Discussion

Our previous study demonstrated that heterotrimeric G proteins and non-canonical Gβ components positively regulate the virulence and secondary metabolism (Yan and Shim 2020). However, in addition to these core components, RGS proteins are well known as negative regulators of G protein signaling pathway to orchestrate this intricate cellular signal transduction mechanism. A further investigation showed that transcription levels of four putative RGS genes *FvFLBA1, FvFLBA2, FvRGSB* and *FvRGSC1* were significantly altered in ΔFvgbb1 deletion mutant when compared to the wild-type progenitor (Mukherjee, et al. 2011). Further characterizations of *F. verticillioides* FvFlbA1 and FvFlbA2, the two FvFlbA paralogs, showed FvFlbA2 is negatively associated with FB1 production (Mukherjee, et al. 2011). Both *FvFLBA1* and *FvFLBA2* mRNA transcription levels are relatively low (data not shown), which suggests that there could be rapid degradation of mRNA after regulating Gα subunits or that the function of these RGS proteins are not transcription dependent. In this study, our aim was to test the hypothesis that two FvFlbA paralogs play unique roles and perhaps complement each other in *F. verticillioides*. Our study revealed that FvFlbA1 function is not critical for FB1 production, whereas the deletion mutants ΔFvflbA2 and ΔFvflbA2/A1 exhibited elevated mycotoxin production and defects in aerial growth.

BrlA is a critical transcription factor involved in activating conidiophore development in both *A. nidulans* and *A. niger*. FlbA was demonstrated to be required for BrlA activation (Lee and Adams 1994). The mutation of *flbA* in *A. nidulans* and *A. niger* resulted in the fluffier hyphal colony (Krijgsheld, et al. 2013). But in *F. verticillioides*, the deletion mutant ΔFvflbA2 showed a less fluffy phenotype similar to that observed in *M. oryzae* ΔMorgs1 whereas ΔFvflbA1 did not exhibit defects in growth (Ramanujam, et al. 2012). ΔFvflbA2 mutant showed reduced conidia production while conidiation was completely hindered in *A. nidulans flbA* mutant. Both ΔFvflbA2 and ΔFvflbA2/A1 deletion mutants showed precocious germination. One possible explanation for this difference in two fungi could be due to distinct asexual production mechanisms between *A. nidulans* and *F. verticillioides*. Conidia in *F. verticillioides* are formed through monophialidic conidiophore that typically develops a single long conidia chain (Leslie and Summerell 2008). *A. nidulans* conidiophore development is drastically different where conidiophore vesicle harbors a layer of metulae and phialides that can each produce a chain of approximately 100 conidia (Yu 2010). *A. nidulans* conidiophore development is one of the most extensively studied models in Ascomycetes. As described earlier, fungal asexual development is proposed to be regulated by Br1A-AbaA-WetA transcription factor cascade (Yu 2010). However, our results in *F. verticillioides* raise the question of whether putative BrlA-AbaA-WetA cascade follows the same expression pattern in two FvFlbA paralogs deletion mutants. Our transcription study showed that expression of these three genes was significantly lower but not completely abolished. This result shows a strong correlation with actual conidia production.

In *A. nidulans*, the *flbA* mutant showed reduced sterigmatocystin production (Hicks, et al. 1997). Interestingly*, A. niger* is known to harbor the putative *FUM* gene cluster and is capable of synthesizing fumonisins (Aerts, et al. 2018). Transcriptome analysis of *flbA* mutant in *A. niger* demonstrated markedly down-regulated expression of *FUM21* which positively corresponds with other *FUM* genes expression and fumonisins production (Aerts, et al. 2018). In *F. verticillioides, FUM21* also functions as a putative Zn(II)_2_Cys_6_ transcription factor that controls expression of genes in the *FUM* cluster, including *FUM1* and *FUM8* (Brown, et al. 2007). *FUM21* locus is located adjacent to *FUM1* that encodes a polyketide synthase responsible for the first step of FB1 production. To further understand how FvFlbA paralogs regulate *FUM* genes expression, we tested transcription levels of *FUM1, FUM8* and *FUM21* in the mutants. Our results showed that these three key *FUM* genes were highly up-regulated in ΔFvflbA2 and ΔFvflbA/A1 mutants, which was consistent with our observation of elevated FB1 production in ΔFvflbA2 and ΔFvflbA2/A1 mutants. The same pattern of FlbA impacts on mycotoxin productions was also reported in the deletion mutants of FlbA paralogs in *F. graminearum* (Park, et al. 2012). FgflbA mutant strain produced significantly higher levels of mycotoxins including DON and ZEA compared to the wild-type progenitor. However, FgFlbB mutant showed no obvious deficiencies in mycotoxin production similar to what we observed in ΔFvflbA1 mutant strain (Park, et al. 2012).

Inactivation of Rgs1 in *M. oryzae* led to precocious appressorium formation on both non-inductive and inductive surfaces, which negatively impacted the pathogen’s ability to cause rice blast disease (Liu, et al. 2007, Zhang, et al. 2011a). Similar outcomes were also observed when DEP or RGS domain were used to complement the ΔMorgs1 mutation that failed to restore virulence on barley and rice (Ramanujam, et al. 2012). This study demonstrated that the DEP or the RGS domain alone could not rescue the infection ability in *M. oryzae*. In addition, *F. graminearum* FgFlbA was shown to play an important role in wheat scab virulence, where ΔFgflbA mutant showing very limited infection limited to inoculated spikelets (Park, et al. 2012). However, our seedling rot assay revealed that FvFlbA1 and FvFlbA2 are not directly associated with virulence in *F. verticillioides*, similar to the Gβ deletion mutant ΔFvgbb1 that did not have influence on stalk rot virulence. This result was consistent with our analyses of stress response and carbon utilization assays, in which both FvflbA1 and FvFlbA2 deletion mutants did not show any defects.

FlbA is well known as a negative regulator of the G protein signaling pathway. One important question we wanted to test was the interactions between two FvFlbA paralogs with canonical G protein components in *F. verticillioides*. Our split luciferase complementation assay demonstrated that FvFlbA1 interact with not only FvFlbA2 but also canonical G protein components. In our experiment, we failed to detect luciferase activity in transformants containing *FvFLBA2-NLuc* with select CLuc constructs. We also performed yeast two-hybrid assays but no interaction between these components were observed (data not shown). With these outcomes, we propose that FvFlbA2 does not directly interact with *F. verticillioides* canonical G protein components, and perhaps we can hypothesize that FvFlbA2 regulates FvFlbA1 and canonical heterotrimeric G protein components indirectly through yet-to-be determined signaling mechanisms. When we tried to test the cellular localization of two FlbA in *F. verticillioides*, we failed to detect the GFP signal in our tested conditions (data not shown). This may be due to the low level of RGS proteins in the cell or that these interactions may be transient under certain developmental or physiological stages of *F. verticillioides*. The mechanisms of how FlbA paralogs are important for FB1 biosynthesis regulation while showing relatively low transcription levels remain unclear. While we predict that two FvFlbA paralogs share same localization with canonical heterotrimeric G protein components, e.g. FvGbb1 and FvGpa2 localized to the cell membranes and vacuoles, respectively (Yan and Shim 2020), further study of FvFlbA cellular functions are necessary to answer these questions.

## Acknowledgements

This research was supported in part by the Agriculture and Food Research Initiative Competitive Grants Program Grant (2013-68004-20359) from the USDA National Institute of Food and Agriculture. The authors declare no conflict of interest.

## Supplementary information

**Fig. S1.**
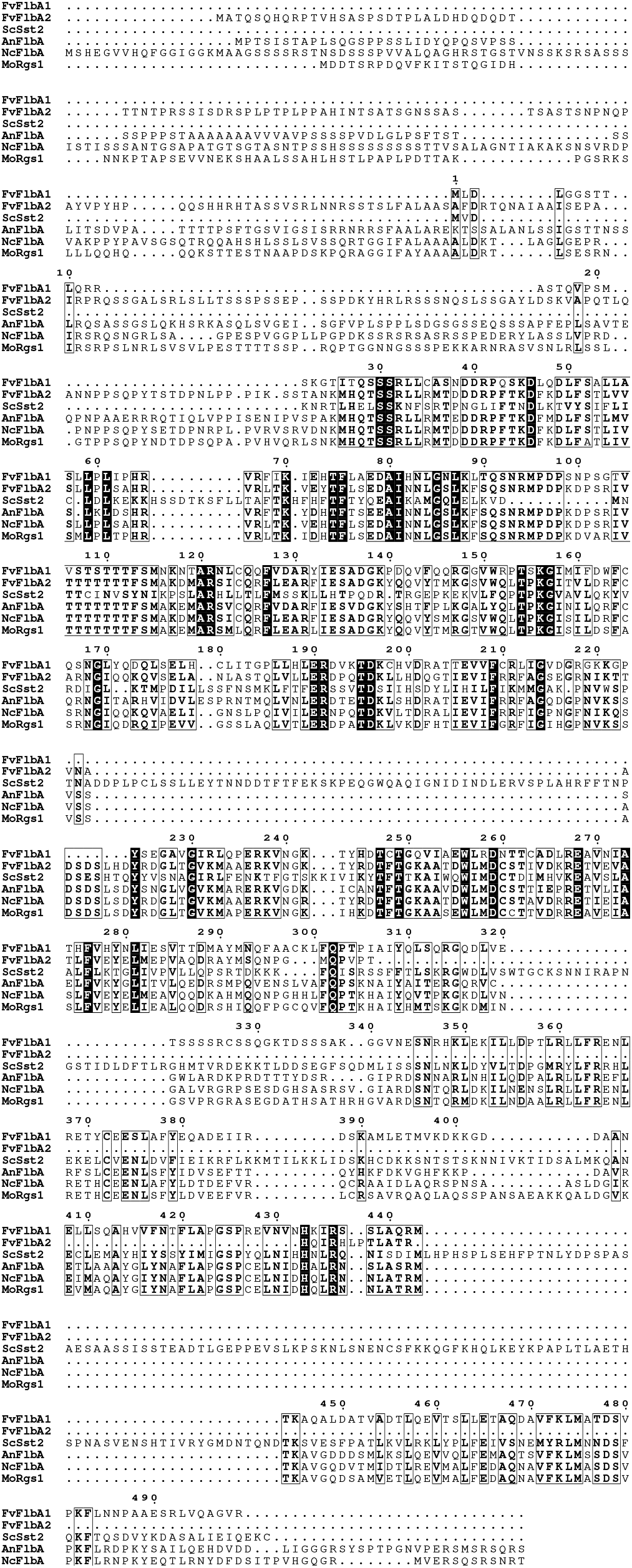
Sequence alignment FlbA ortholog proteins in select fungal species. We aligned protein sequences of *Fusarium verticillioides* FvFlbA1, *F. verticillioides* FvFlbA2, *Saccharomyces cerevisiae* ScSst2, *A.nidulans* AnFlbA, *Neurospora crassa* NcFlbA, and *Magnaporthe oryzae* MoRgs1. Identical and similar sequences were displayed in the box.

**Fig. S2.**
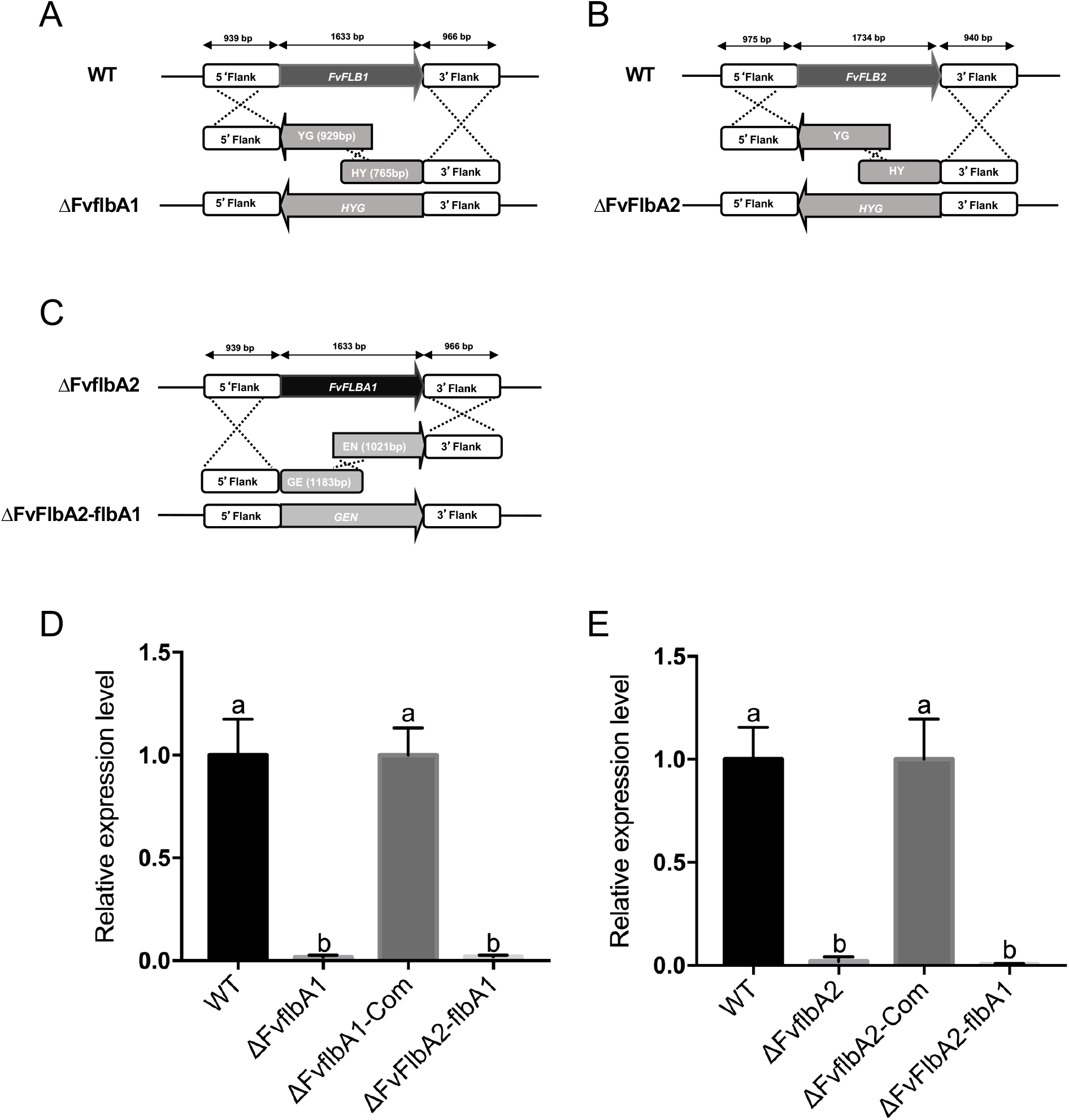
Split marker approach employed in *FLBA* generating gene deletion mutants in *F. verticillioides*. (A) ΔFvflbA1 mutant was generated by replacing *FLBA1* gene with a hygromycin B phosphotransferase gene (*HPH*). (B) *FvFLBA2 gene* was replaced by a hygromycin gene to generate a ΔFvflbA2 mutant. (C) ΔFvflbA2/A1*FvFLBA1* was generated by replacing *FLBA1* gene with a geneticin gene (*GEN*) in ΔFvflbA2 background. (D) Mutants ΔFvflbA1, ΔFvflbA1-Com, ΔFvflbA2/A1 were subject to qPCR using *FvFLBA1* gene primer. Relative expression was normalized to *F. verticillioides* β-tubulin gene (FVEG_04081). (E) *FvFLBA2* gene transcription level was examined in WT, ΔFvflbA2, ΔFvflbA2-Com, ΔFvflbA2/A1 strains.

**Fig. S3.**
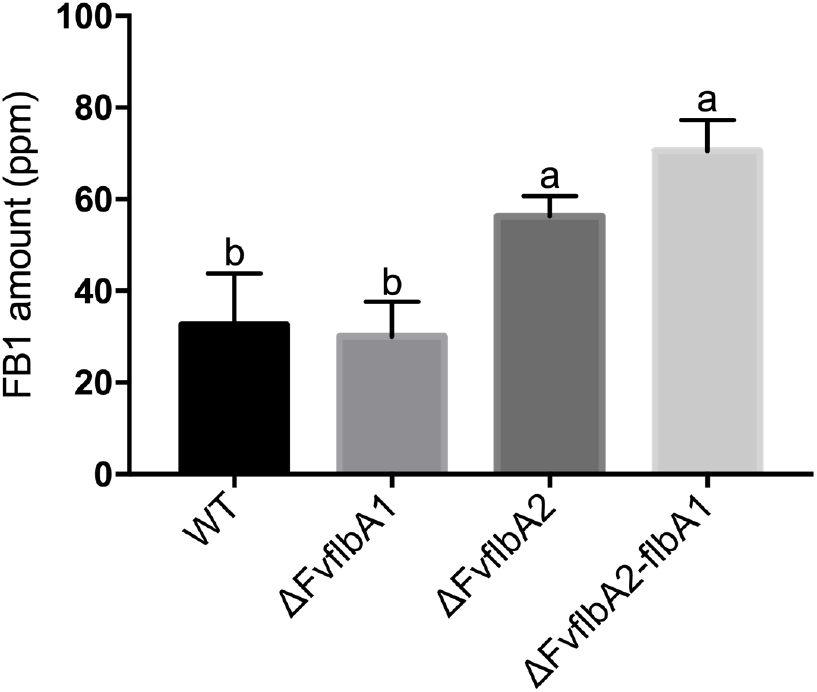
Relative FB1 expression in wild-type and FvflbA mutants. FB1 was extracted from supernatant (2ml) of 7-day incubation at 150 RPM in myro liquid culture. In details, YEPD mycelial samples (3 dpi) were harvested through Miracloth (EMD Millipore). Subsequently, mycelia (0.3g) were inoculated into 100 ml myro liquid medium at room temperature.

